# The hypoxic ventilatory response is facilitated by the activation of Lkb1-AMPK signalling pathways downstream of the carotid bodies

**DOI:** 10.1101/604900

**Authors:** Amira D. Mahmoud, Andrew P. Holmes, Sandy MacMillan, Oluseye A. Ogunbayo, Christopher N. Wyatt, Mark L. Dallas, Prem Kumar, Marc Foretz, Benoit Viollet, A. Mark Evans

## Abstract

We recently demonstrated that the role of the AMP-activated protein kinase (**AMPK**), a ubiquitously expressed enzyme that governs cell-autonomous metabolic homeostasis, has been extended to system-level control of breathing and thus oxygen and energy (ATP) supply to the body. Here we assess the contribution to the hypoxic ventilatory response (HVR) of two upstream kinases that govern the activities of AMPK. Lkb1, which activates AMPK in response to metabolic stress and CaMKK2 which mediates the alternative Ca^2+^-dependent mechanism of AMPK activation. HVRs remained unaffected in mice with global deletion of the CaMKK2 gene. By contrast, HVRs were markedly attenuated in mice with conditional deletion of LKB1 in catecholaminergic cells, including carotid body type I cells and brainstem respiratory networks. In these mice hypoxia evoked hypoventilation, apnoea and Cheyne-Stokes-like breathing, rather than hyperventilation. Attenuation of HVRs, albeit less severe, was also conferred in mice carrying ∼90% knockdown of Lkb1 expression. Carotid body afferent input responses were retained following either ∼90% knockdown of Lkb1 or AMPK deletion. In marked contrast, LKB1 deletion virtually abolished carotid body afferent discharge during normoxia, hypoxia and hypercapnia. We conclude that Lkb1 and AMPK, but not CaMKK2, facilitate HVRs at a site downstream of the carotid bodies.

## 1. BACKGROUND

The AMP-activated protein kinase (AMPK) is a cellular energy sensor that maintains cell-autonomous energy homeostasis. From its 2 α (catalytic), 2 β and 3 γ (regulatory) subunits 12 AMPK heterotrimers may be formed, each harbouring different sensitivities to activation by increases in cellular AMP and ADP, and the capacity to directly phosphorylate and thus regulate different targets [1]. AMPK is coupled to mitochondrial oxidative phosphorylation by two discrete albeit cooperative pathways, involving liver kinase B1 (Lkb1) and changes in the cellular AMP:ATP and ADP:ATP ratios. Binding of AMP to the AMPK γ subunit increases activity 10-fold by allosteric activation alone, while AMP or ADP binding delivers increases in Lkb1-dependent phosphorylation and reductions in dephosphorylation of Thr172 on the α subunit that confer 100-fold further activation. All of these effects are inhibited by ATP [2]. Lkb1 is, therefore, the principal pathway for AMPK activation during metabolic stresses such as hypoxia. However, there are alternative Ca^2+^-dependent pathways to AMPK activation that are governed by the calmodulin-dependent protein kinase CaMKK2, which delivers increases in Thr172 phosphorylation and thus AMPK activation independent of changes in cellular AM(D)P:ATP ratios.

Classically AMPK regulates cell-autonomous pathways of energy supply by phosphorylating targets that switch off non-essential anabolic processes that consume ATP and switch on catabolic pathways that generate ATP, thereby compensating for deficits in ATP supply or availability[1]. Recently, however, we demonstrated [3] that the role of AMPK in metabolic homeostasis is not limited to such cell autonomous pathways, but extends to the hypoxic ventilatory response (HVR)[4, 5] and thus O_2_ and energy (ATP) supply to the body as a whole. In doing so AMPK acts to oppose central respiratory depression during hypoxia and thus resists hypoventilation and apnoea. Surprisingly, however, AMPK deficiency did not precipitate ventilatory dysfunction at the level of the carotid bodies as one would predict given their role as the primary arterial chemoreceptors [5-8], but attenuated activation during hypoxia of the caudal brainstem while afferent input responses from the carotid bodies were normal [3]. We therefore hypothesised that AMPK may aid delivery of HVRs by integrating “local hypoxic stress” within brainstem respiratory networks with an index of “peripheral hypoxic status” provided via afferent chemosensory inputs. In this respect it is clear that the capacity for signal integration could be determined centrally either through AMPK activation consequent to brainstem hypoxia, increases in the AM(D)P:ATP ratio and thus Lkb1-dependent phosphorylation, or CaMKK2-dependent phosphorylation in response to increases in cytoplasmic Ca^2+^.

Using a variety of genetic mouse models, along with both *in vitro* and *in vivo* approaches, the present study examines whether Lkb1 and/or CaMKK2 are important drivers of HVRs. We reveal that Lkb1-AMPK signalling pathways facilitate HVRs independent of CaMKK2.

## 2. RESULTS

Because global gene deletion of *Lkb1* or *AMPKα1+α2* is embryonic lethal we employed conditional deletion of these genes. For *Lkb1* deletion we used floxed mice in which the gene enoding Lkb1 (*Stk11*) had been replaced by a cDNA cassette encoding equivalent exon sequences, and exon 4 and the cDNA cassette flanked by loxP sequences, which in their own right deliver ∼90% global knockdown of Lkb1 expression [9]. For *AMPKα1+α2* deletion critical exons of the genes encoding AMPKα1 (*Prkaa1*) and AMPKα2 subunits (*Prkaa2*) were flanked by loxP sequences [10]. Each floxed mouse line was crossed, as previously described [3], with mice expressing Cre recombinase under the control of the tyrosine hydroxylase (TH) promoter, providing for gene deletion in all catecholaminergic cells inclusive of those cells that constitute the hypoxia-responsive respiratory network from carotid body [8] to brainstem [11]. Transient developmental expression of TH does occur in disparate cell types that do not express TH in the adult [12], such as dorsal root ganglion cells and pancreatic islets, but these do not contribute to the acute HVR. We previously confirmed restriction of Cre to TH-positive cells in the adult mouse by viral transfection of a Cre-inducible vector carrying a reporter gene[3]. Therefore, our approach overcomes embryonic lethality and allows, unforeseen ectopic Cre expression aside, for greater discrimination of circuit mechanisms than would be provided for by global knockouts. The role of CaMKK2 in the hypoxic ventilatory response (HVR) was determined by assessing mice with global deletion of the corresponding gene (*CaMKK2*) [13].

Under normoxia there was no difference between controls and either *Lkb1, CaMKK2* or, as previously shown, *AMPKα1+α2* knockouts[14] with respect to weight versus age, breathing frequency, tidal volume or minute ventilation (**Supplementary Fig 1 and 2**). Nevertheless, profound and genotype-specific differences were observed with respect to the ventilatory responses during hypoxia and hypercapnia.

**Figure 1.**
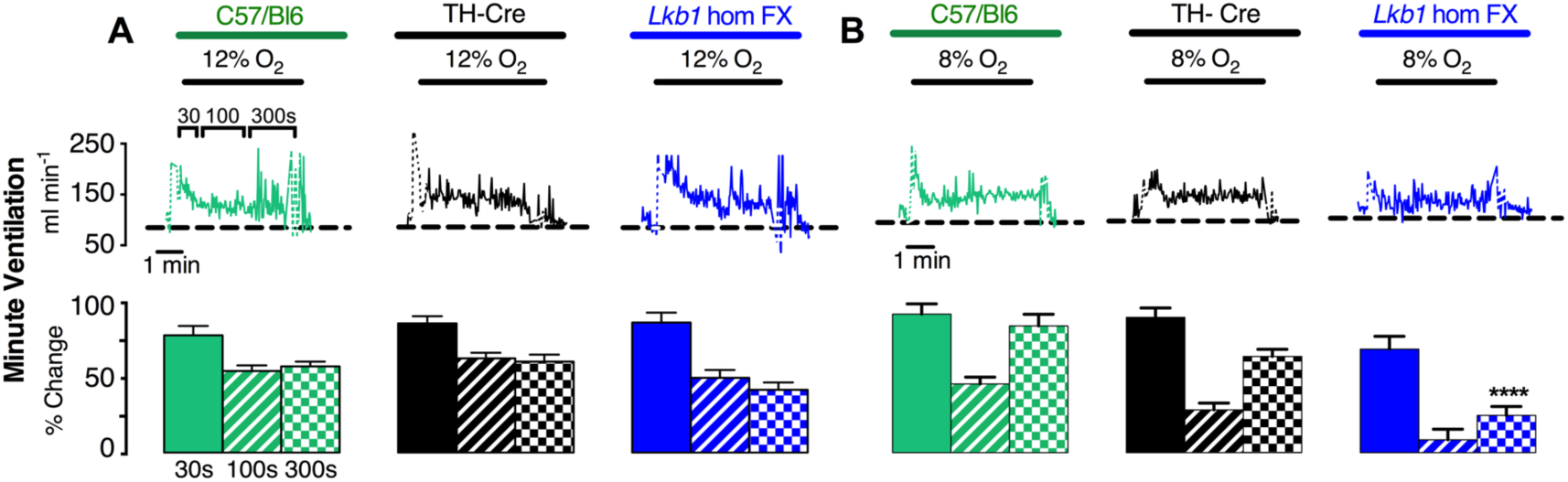
Mice hypomorphic for Lkb1 exhibit an attenuated hypoxic ventilatory response measured by unrestrained plethysmography. *Upper panels* show example records and *lower panels* bar charts of mean±SEM for increases in minute ventilation at the peak of the Augmenting Phase (A, ∼30s), after Roll Off (RO, ∼100s) and during the plateau of the Sustained Phase (SP, ∼300s) of the ventilatory response to (*A*) 12% and (*B*) 8% O_2_ for C57Bl6 (green, n = 6), TH-Cre (black, n = 31) and *Lkb1* homozygous floxed mice (*Lkb1* hom FX, blue, n = 14) which are globally hypomorphic (∼90% loss of Lkb1). ****=p< 0.0001 relative to TH-Cre and C57/Bl6.

**Figure 2.**
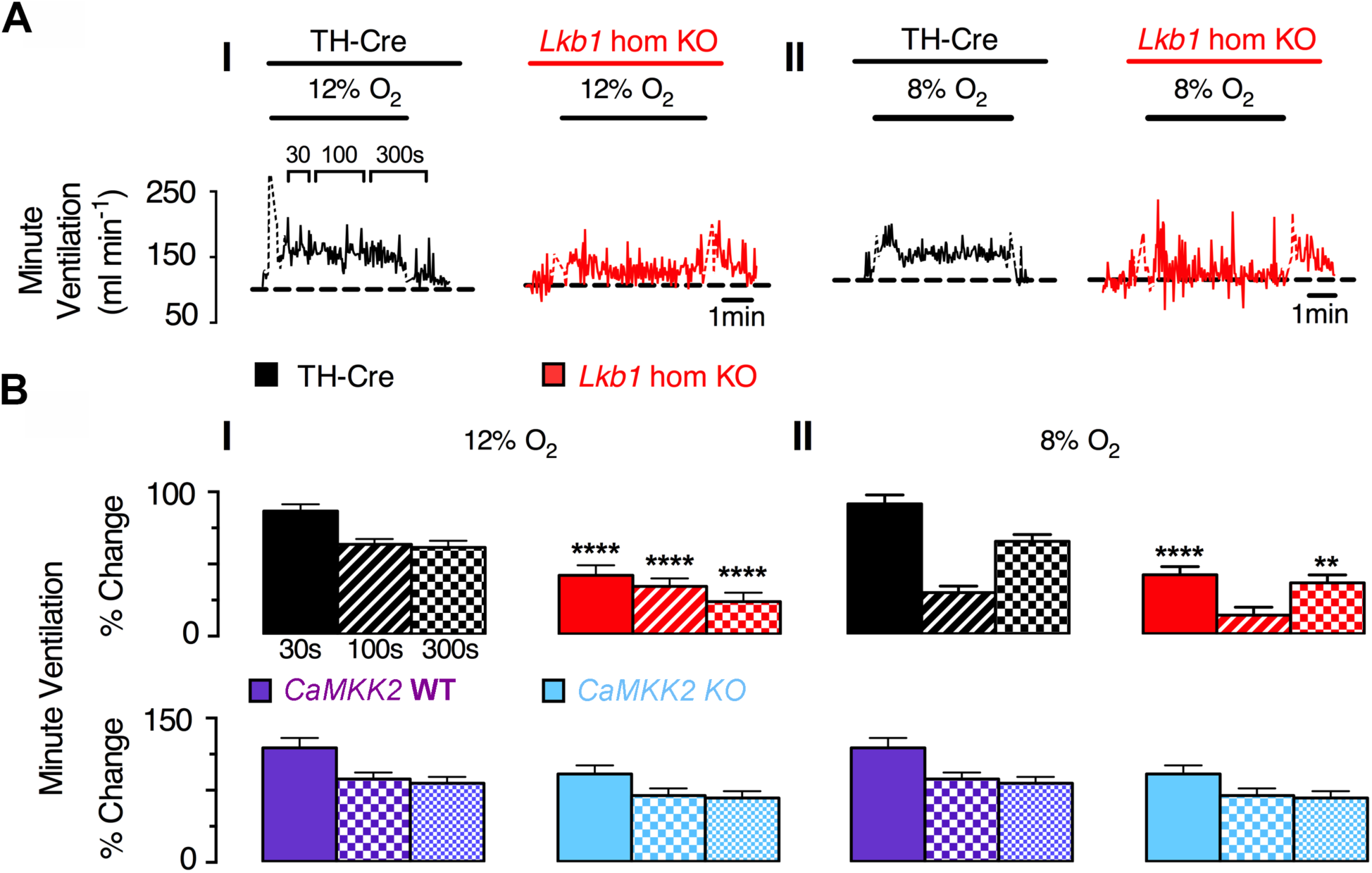
Conditional deletion of *Lkb1* in tyrosine hydroxylase expressing cells markedly attenuates the hypoxic ventilatory response measured by unrestrained plethysmography, but global *CaMKK2* knockout does not. A, Example records of minute ventilation (ml min^−1^ g^−1^) versus time during (I) 12% O_2_ and (II) 8% O_2_ for TH-Cre (black, n = 31) and conditional *Lkb1* knockout mice (*Lkb1* hom KO, red, n = 22). B, Bar charts show mean±SEM for changes in minute ventilation at the peak of the Augmenting Phase (A, ∼30s), at 100s following Roll Off (RO, ∼100s) and the plateau of the Sustained Phase (SP, 300s) of the response to hypoxia of (*Upper Panels*) TH-Cre (black, n = 31) conditional *Lkb1* hom KO (red, n = 22) and (*Lower Panels*) *CaMKk2 WT* (purple, n = 7) and global *CaMKK2* KO (blue, n = 7). *=p<0.05, **** = p < 0.00001.

### 2.1 HVRs are attenuated in mice with Lkb1 deficiency but remain unaffected by global CaMKK2 deletion

Attenuation of HVRs was observed in *Lkb1* floxed mice used here, which harbour ∼90% global Lkb1 deficiency [9]. In this respect it is notable (see below) that this effect only reached significance, compared to controls (C57/Bl6 and TH-Cre), during the sustained phase of the response to severe but not moderate hypoxia (**Fig 1A-B**), i.e. these mice exhibited delayed hypoventilation during hypoxia. This provides indirect support of our proposal that AMPK facilitates HVRs downstream of the carotid bodies, because this observation is accordance with the view that carotid body chemoafferent input responses drive the augmenting phase of HVRs [15, 16] while activation by hypoxia of brainstem respiratory networks may provide for maintenance of HVRs in the longer term [3, 5, 16-19].

The effect of conditional *Lkb1* deletion (**Fig 2, and Supplementary Fig 3**) was more severe, with HVRs suppressed, relative to controls (TH-Cre), during 5 min exposures to either mild (12% O_2_; **Fig 2AI and BI *upper panels***) or severe hypoxia (8% O_2_; **Fig 2AII and BII *upper panels***) and in a manner proportional to the severity of hypoxia. Relative to control mice (TH-Cre), the peak change in minute ventilation (∼30s) achieved by *Lkb1* knockouts during the initial “Augmenting Phase” was lower (P<0.0001, compared to TH-Cre). This is important because there is general agreement that this phase of the HVR primarily results from carotid body afferent input responses [4, 5, 15]. Minute ventilation was similarly depressed relative to controls following subsequent ventilatory depression (Roll Off, ∼100s, NS) and during the latter sustained phase of the response to hypoxia (2-5min; P<0.0001). Note: 0.05% CO_2_ used here is probably insufficient to prevent significant respiratory alkalosis which may have impacted ventilatory reflexes during the latter phases of the sustained hypoxic stimulus [20] of wild type mice in particular. Therefore, we may have underestimated the degree to which *Lkb1* deletion inhibits HVRs.

**Figure 3.**
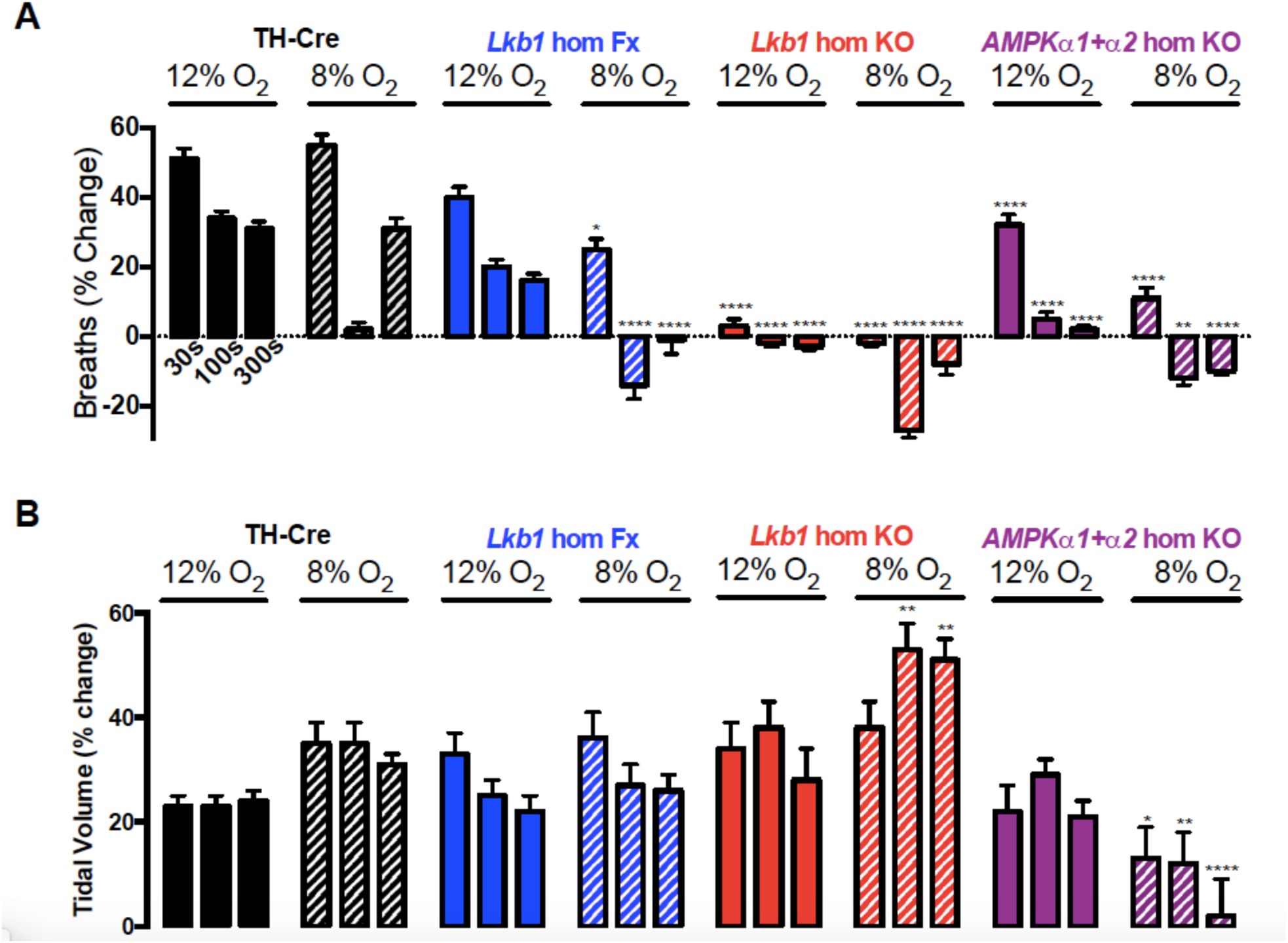
Conditional deletion of *Lkb1* in tyrosine hydroxylase expressing cells blocks increases in breathing frequency but augments increases in tidal volume during severe hypoxia measured by unrestrained plethysmography. Bar charts of mean±SEM for increases in (A) Breathing frequency and (B) Tidal volume At the peak of the Augmenting Phase (∼30s), at ∼100s following Roll Off and during the plateau of the Sustained Phase (∼300s) of the ventilatory response to hypoxia for TH-Cre (black, n = 31), *Lkb1* homozygous floxed mice (*Lkb1* hom Fx, blue, n = 22) that are ∼90% hypomorphic for Lkb1 and conditional *Lkb1* homozygous knockout mice (*Lkb1* hom KO, red, n = 22). These data are also compared with outcomes for conditional *AMPKα1*+*α2* homozygous knockout mice (*AMPKα1*+*α2* hom KO, mauve, n = 26-30). *=p<0.05, **=p<0.01, ****=p< 0.0001 compared to TH-Cre.

By contrast, global deletion of CaMKK2 did not affect HVRs in any discernable way (**Fig 2BI-II *lower panels***), ruling out a prominent role for AMPK activation through this alternative Ca^2+^-dependent pathway.

### 2.2 Attenuation of HVRs by *Lkb1* deletion results from deficits in breathing frequency

Intriguingly, deficits in minute ventilation in *Lkb1* floxed mice resulted from attenuation of increases in breathing frequency at all time points during exposure to severe (8% O_2_; n=22; P≤0.0001) but not mild (12%; n=15; NS) hypoxia. Increases in breathing frequency were even more markedly attenuated by homozygous *Lkb1* deletion and at all time points during exposure to both mild and severe hypoxia when compared to controls (TH-Cre; P<0.0001; **Fig 3A**), albeit in a manner proportional to the severity of hypoxia. By contrast no attenuation of increases in tidal volume were observed for either *Lkb1* floxed mice or *Lkb1* knockouts during mild or severe hypoxia.

The aforementioned findings are intriguing because it has been proposed that peripheral chemoreceptors primarily drive increases in breathing frequency during moderate to severe hypoxia (see for example [21]). It is therefore all the more important to note that mice with conditional deletion of *AMPKα1+α2* in catecholaminergic cells, which retain carotid body afferent input responses, exhibit markedly attenuated increases in breathing frequency but not tidal volume at all time points when exposed to mild hypoxia (12% O_2;_ **Fig 3A-B**). By contrast, however, *AMPKα1+α2* deletion attenuated increases in both breathing frequency and tidal volume during severe (8% O_2_) hypoxia [14]. When taken together these findings strongly suggest that Lkb1-AMPK signalling pathways facilitate HVRs and oppose respiratory depression during hypoxia. Moreover, outcomes suggest that those circuit mechanisms that mediate hypoxia-evoked increases in tidal volume are afforded greater protection from the impact of Lkb1 and AMPK deficiency than those delivering increases in breathing frequency.

### 2.3 Lkb1 deficiency causes ventilatory instability and Cheyne-Stokes-like breathing during hypoxia

Quite unlike our previously reported findings in mice with *AMPKα1+α2* deletion[14], average measures (excluding apnoeas) for *Lkb1* knockouts indicated significant augmentation rather than attenuation of increases in tidal volume during severe hypoxia, relative to controls (P<0.01 at 100s and 300s compared to TH-Cre). Closer inspection revealed that attenuation of HVRs in *Lkb1* knockouts during exposure to 8% O_2_ only, was associated with periods of Cheyne-Stokes-like breathing (CSB), i.e., tidal volume exhibited marked, sinusoidal variations with time (**Fig 4A and B**). Periods of CSB in *Lkb1* knockout mice were generally separated by frequent, prolonged apnoeas (≤4s), with apnoea frequency, apnoea duration and apnoea duration index (frequency × duration) all significantly larger (P<0.0001) than for controls (**Fig 4C**). Nevertheless, as might be expected given outcomes for minute ventilation, apnoea frequency and duration also increased in a manner directly related to the severity of hypoxia. Moreover, CSB and increases in apneoa frequency and duration observed during severe hypoxia were completely reversed by hypercapnic hypoxia (**Fig 4AIII and C**), likely due to improved O_2_ supply consequent to increases in ventilation (see below). The appearance of CSB likely accounts for measured increases in tidal volume for these mice relative to controls. That aside it is important to note that periods of hypoxia-evoked CSB in *Lkb1* knockouts occurred irrespective of whether they were preceded by spontaneous or post-sigh apnoeas (**Fig 4B**). Moreover both spontaneous and post-sigh apnoeas were equally frequent during exposure of *AMPKα1+α2* knockouts to severe hypoxia, where CSB is absent during 5 min [3] or even 10 min (**Fig 5**) exposures to severe hypoxia. In short, if sighs are triggered by hypoxia at a given threshold [3, 22], central hypoxia is likely no more severe for *Lkb1* when compared to *AMPKα1+α2* knockouts and CSB consequent to Lkb1 deficiency is thus most likely triggered by other means.

**Figure 4.**
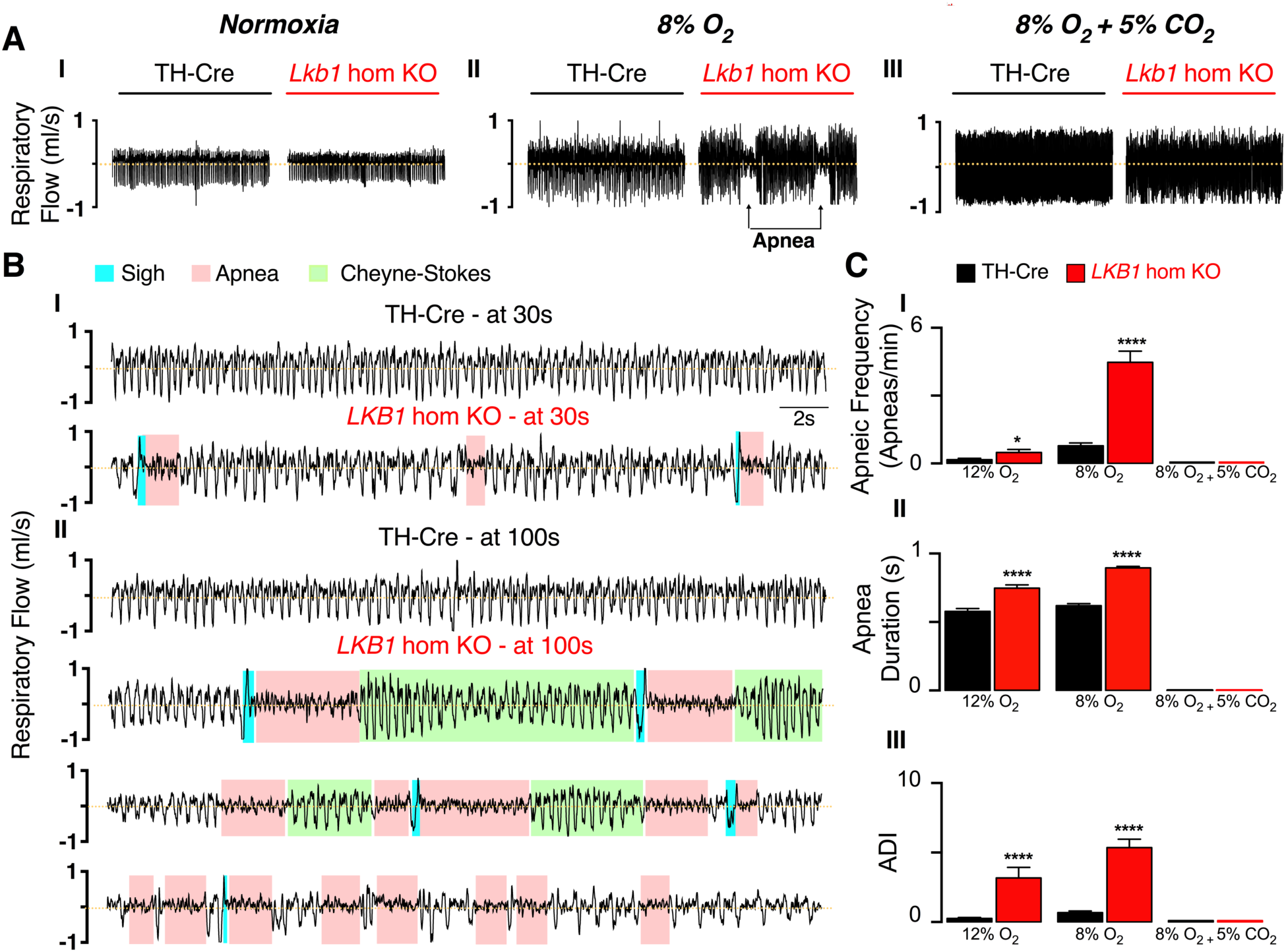
Unrestrained plethysmography shows that conditional deletion of *Lkb1* in tyrosine hydroxylase expressing cells precipitates hypoventilation, apnoea and Cheyne-Stokes-like breathing during severe hypoxia. A, Example records of ventilatory activity from TH-Cre and conditional *Lkb1* homozygous knockout mice (*Lkb1* hom KO) during (*I*) normoxia (21% O_2_), (*II*) hypoxia (8% O_2_) and (*III*) hypoxia with hypercapnia (8% O_2_ + 5% CO_2_), that were obtained using whole body plethysmography. B(I-II), Typical ventilatory records for TH-Cre and conditional *Lkb1* hom KO mice on an expanded time scale at the indicated time points during exposures to severe hypoxia (8% O_2_). C, mean±SEM for (*I*) apnoeic index (per minute), (*II*) apnoea duration (s) and (*III*) apnoea-duration index (frequency × duration) for TH-Cre (black, n = 31) and conditional *Lkb1* hom KO (red, n = 22) mice during exposures to 12% O_2_, 8% O_2_ and 8% O_2_ + 5% CO_2_. *=p<0.05, ****=p< 0.0001.

**Figure 5.**
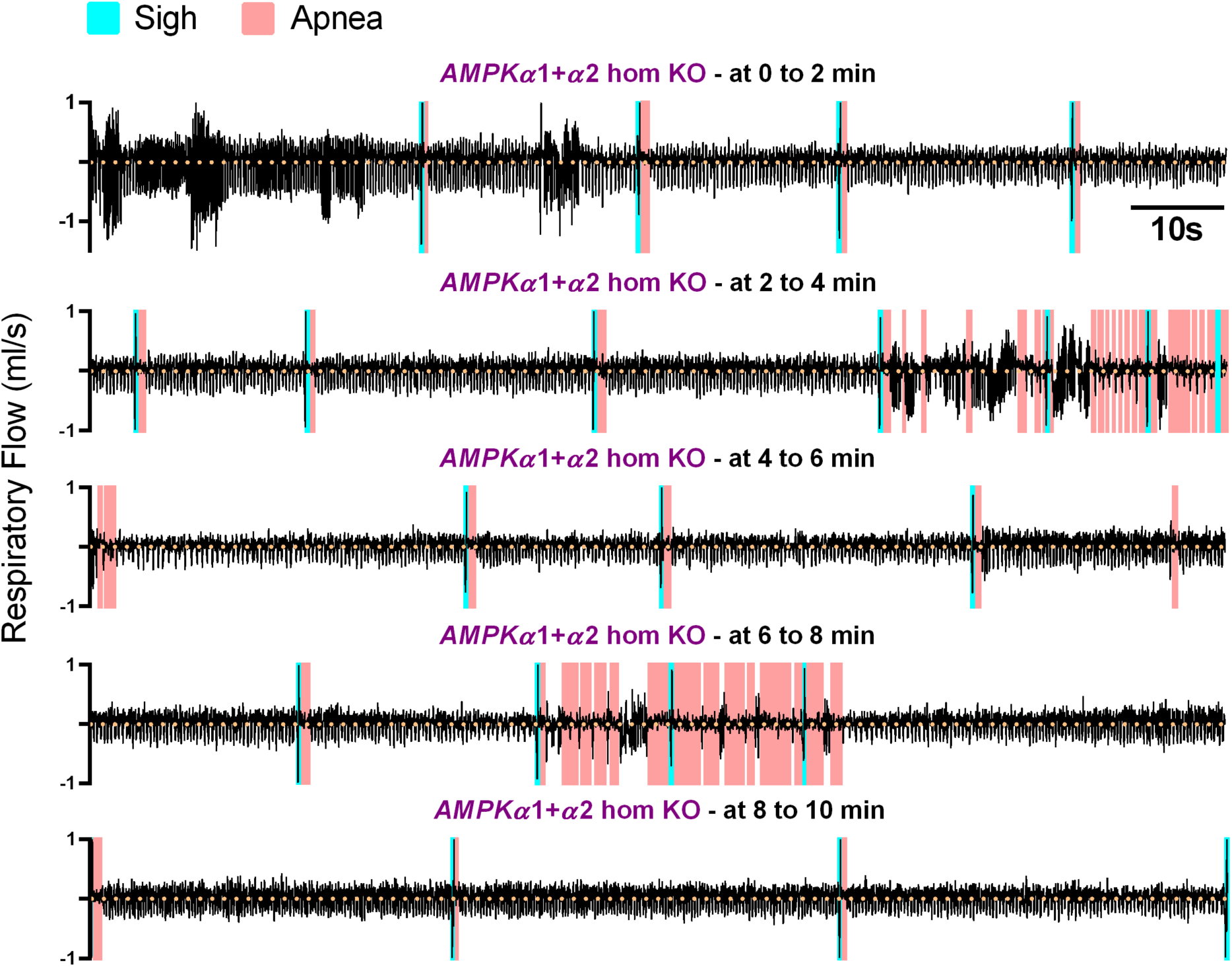
Conditional deletion of *AMPK-α1+α2* in tyrosine hydroxylase expressing cells does not precipitate Cheyne-Stokes-like ventilation even during prolonged 10 minute exposures to severe hypoxia during unrestrained plethysmography. Typical ventilatory record for conditional *AMPK-α1+α2* double knockout mice (*AMPKα1+α2* hom KO) during a 10 minute exposure to severe hypoxia (8% O_2_).

### 2.2 Conditional *Lkb1* deletion slows the ventilatory response to hypercapnic hypoxia and hypercapnia

The ventilatory response to hypercapnic hypoxia (8% O_2_ + 5% CO_2_) remained entirely unaffected following *AMPKα1+α2* deletion [3]. By contrast, in *Lkb1* knockouts increases in minute ventilation were attenuated, but only during the rising phase of the ventilatory response to hypercapnic hypoxia (P<0.01; **Fig 6A**), indicating that *Lkb1* deletion slowed the rising phase of the response to this stimulus but did not affect the peak achieved. It is possible that this may reflect the partial restoration of the initial rise in respiratory frequency during hypercapnic hypoxia, that is attenuated during hypoxia alone (**Supplementary Fig 4**). However, the rise in minute ventilation during exposure to hypercapnia alone (5% CO_2_) was also markedly slower for *Lkb1* knockouts (P<0.05), but thereafter achieved an equivalent magnitude (**Fig 6B**), as a consequence of equivalent peak increases in both respiratory frequency and tidal volume **(Supplementary Fig 4**). By contrast, mice with *AMPKα1+α2* deletion had preserved peak hypercapnic ventilatory responses without any initial delay in onset (**Fig 6A-B**).

**Figure 6.**
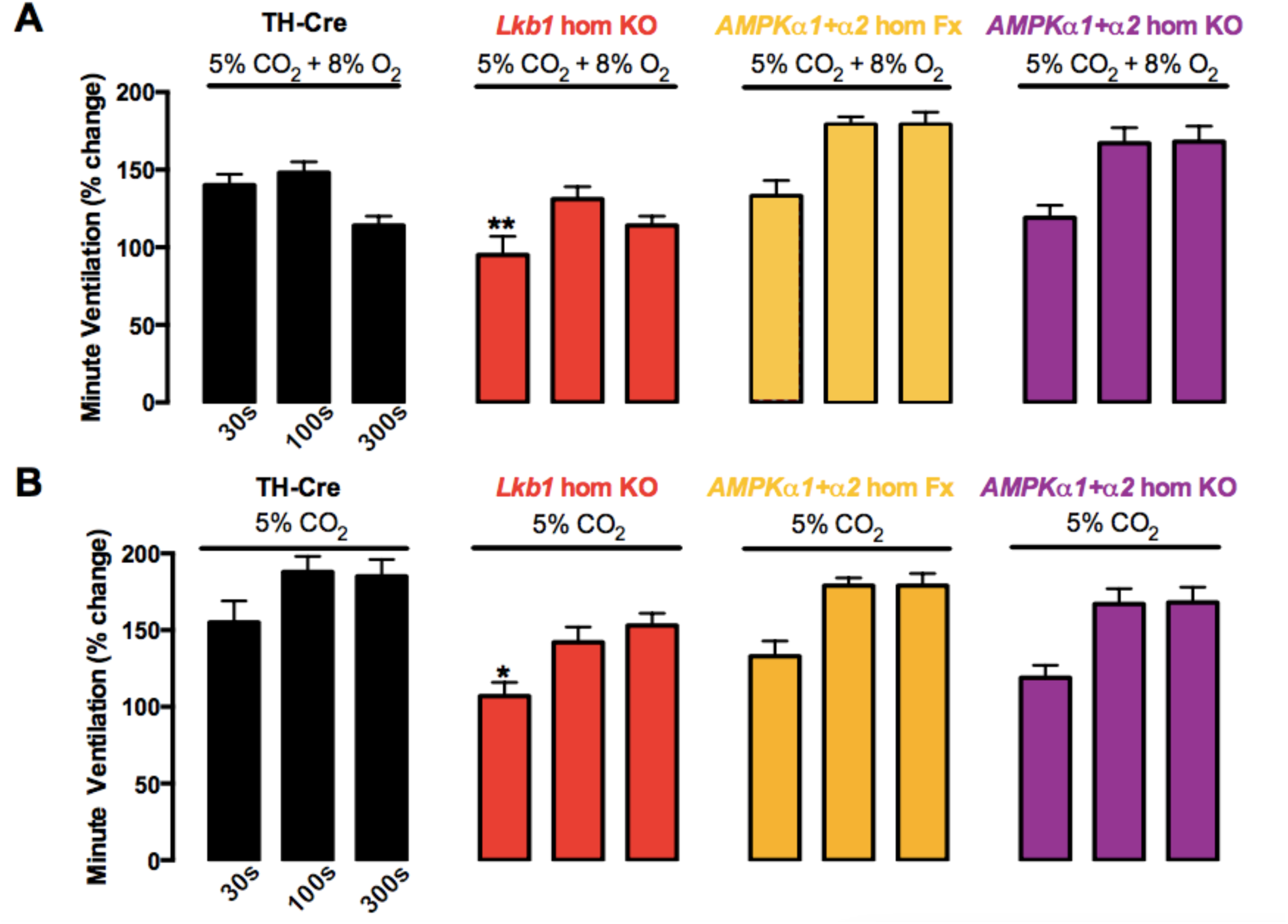
Conditional deletion of Lkb1 in tyrosine hydroxylase expressing cells markedly slows the hypercapnic ventilatory response measured by unrestrained plethysmography. Bar charts of mean±SEM for increases in minute ventilation at ∼30s, 100s and 300s during exposures to (A) Hypercapnic hypoxia (5% CO_2_+8%O2) and (B) Hypercapnia (5% CO_2_) for TH-Cre (black, n = 31), conditional *Lkb1* homozygous knockout mice (*Lkb1* hom KO, red, n = 22), *AMPKα1*+*α2* homozygous floxed mice (*AMPKα1*+*α2* hom Fx, beige, n = 26) and *AMPKα1*+*α2* homozygous knockout mice (*AMPKα1*+*α2* hom KO, mauve, n = 30). *=p<0.05, **=p< 0.0001.

### 2.3 *Lkb1* but not *AMPKα1+α2* deletion attenuates carotid body chemoafferent discharge during normoxia, hypoxia and hypercapnia

During normoxia mean±SEM basal afferent fibre discharge frequency from *in-vitro* carotid bodies of TH-Cre mice was similar to that for carotid bodies of homozygous *Lkb1* floxed mice that harbour ∼90% global Lkb1 deficiency [9]. In marked contrast, however, basal afferent discharge measured from carotid bodies of *Lkb1* knockouts was markedly attenuated (**Fig 7C**; n=7; P≤0.001 versus TH-Cre and *Lkb1* floxed).

**Figure 7.**
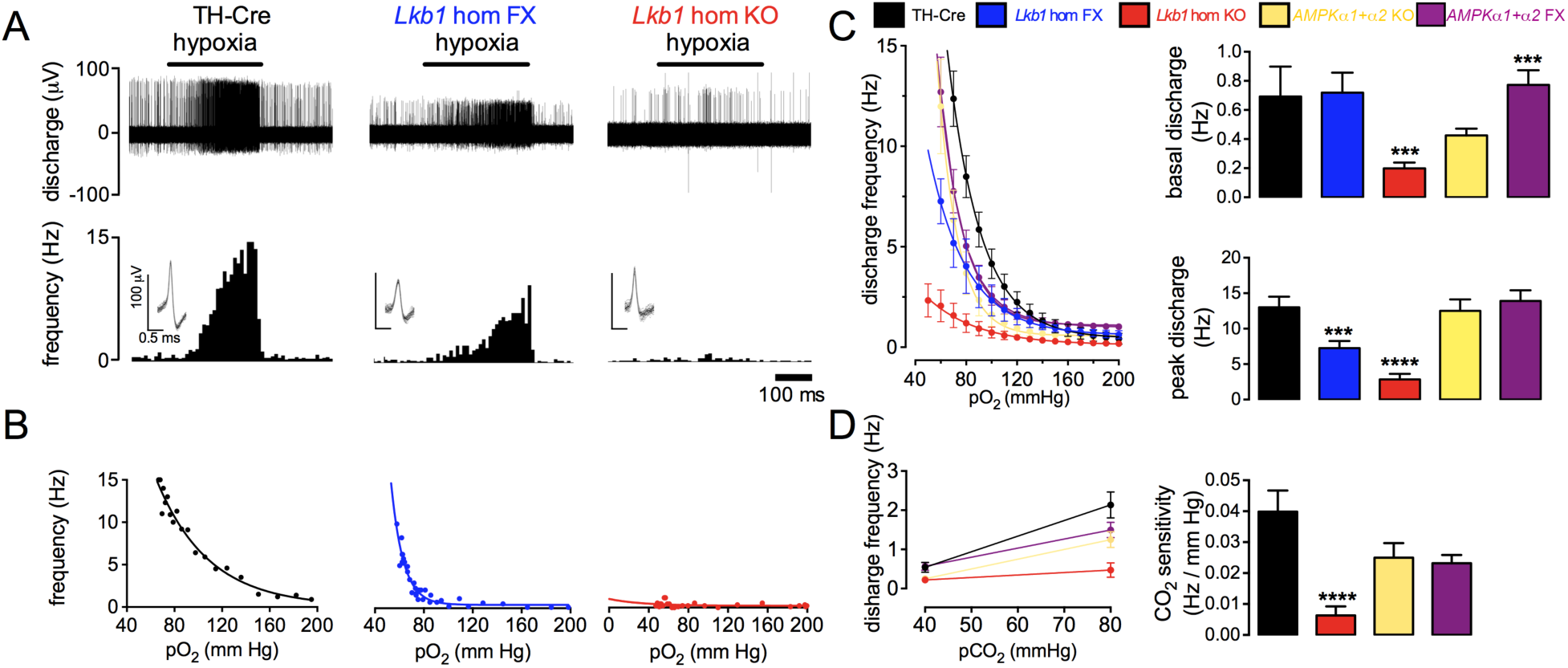
Conditional deletion of *Lkb1* but not *AMPK-*α*1+*α2 in tyrosine hydroxylase positive cells attenuates afferent discharge from the carotid body in-vitro during normoxia, hypoxia and hypercapnia. A, *Upper panels* show extracellular recordings of chemoafferent discharge versus time for carotid bodies from control (TH-Cre, black), *Lkb1* homozygoous floxed (*Lkb1* hom FX, blue) and conditional *Lkb1* homozygous knockout (*Lkb1* hom KO, red) mice, *Lower panels* show frequency-time histograms (*inset:* single fibre discriminations). B, Frequency-*p*O_2_ response curves corresponding to records in (*A*). C, *Left hand panel* compares mean±SEM for frequency-*p*O_2_ response curves for TH-Cre (black, n = 8), *Lkb1* hom FX (blue, n = 7), conditional *Lkb1* hom KO (red, n = 7), *AMPK-*α*1* and -α2 double floxed *(AMPK* hom FX, beige, n=5) and conditional *AMPK-α1*+α2 double knockouts (*AMPK* hom DKO, mauve, n=5). Bar charts in *right hand panels* show mean±SEM for (*upper*) basal single fibre discharge frequency and (*lower*) peak single fibre discharge frequency during hypoxia. D, Mean±SEM for chemoafferent discharge versus *pC*O_2_ (*left hand panel*) and CO_2_ sensitivity (*right hand panel*) for TH-Cre, *Lkb1* hom KO, *AMPK* hom FX *and AMPK* hom DKO. *** =p<0.001, ****=p< 0.0001.

In line with the above, reductions in superfusate *P*O_2_ increased chemoafferent discharge from carotid bodies of TH-Cre mice exponentially, but evoked only marginal increases in afferent discharge from carotid bodies of *Lkb1* knockouts (P<0.0001; **Fig 7C**). By contrast, peak discharge frequencies (*P*O_2_ ≤75 mmHg) of carotid bodies from hypomorphic *Lkb1* floxed mice were attenuated by less than 50% relative to controls, despite the fact that these mice exhibit global deficits in Lkb1 expression of ∼90% [9]

For *AMPKα1+α2* deletion basal afferent discharge frequency was higher than recorded for *AMPKα1+α2* floxed mice (n=5; P<0.001) [3], but not significantly different from measures for either TH-Cre (n = 8) or the hypomorphic *Lkb1* floxed (n =7) mice. *AMPKα1+α2* floxed mice may represent the better comparison, raising the possibility that AMPK may ordinarily act to reduce basal afferent discharge frequency through, for example, inhibition of the large conductance voltage- and calcium-activated potassium current (BK_Ca_)[23]. However, it is clear that this does not hold when basal discharge of *AMPKα1+α2* knockouts is compared against our full range of control mice. Moreover, peak discharge frequency during hypoxia (**Fig 7C**) was similar for *AMPKα1+α2* knockouts when compared to *AMPKα1+α2* floxed (n=5; NS) and TH-Cre mice (n=8; NS).

Carotid bodies isolated from *AMPKα1+α2* knockouts also had a preserved chemoafferent response to hypercapnia. By contrast, *Lkb1* deletion (n=4) inhibited carotid body responses to hypercapnia and reduced carotid body CO_2_-sensitivity (which is linear between 40 and 80 mmHg) when compared to TH-Cre (n=7; **Fig 7D**). This may explain, in part, the slower rising phase of the ventilatory response of *Lkb1* knockouts during hypercapnic hypoxia and hypercapnia, as carotid body afferent inputs to the brainstem determine this [15, 24].

### 2.4 The rank order of severity for HVR versus carotid body dysfunction is different

As demonstrated above, while *Lkb1* deletion virtually abolished, hypomorphic Lkb1 expression modestly attenuated and *AMPKα1+α2* deletion was without effect on afferent output from the carotid body during hypoxia, all three interventions markedly attenuated the HVR during severe hypoxia. Moreover, the order of severity for hypoxic ventilatory dysfunction was different when compared to that for inhibition of carotid body chemoafferent discharge during hypoxia. This is evident from **Fig 8**, which compares Poincaré plots of inter-breath interval (BBn) versus subsequent inter-breath interval (BBn+1) during normoxia (**AI**), 12% O_2_ (**BI**), 8% O_2_ (**CI**), 8% O_2_ + 5% CO_2_ (**DI**) for controls (TH-Cre), hypomorphic *Lkb1* floxed mice and *Lkb1* knockouts (**Fig 8AI-DI**, *left hand panels*) with previously published [3] examples for *AMPKα1+α2* knockouts (**Fig 8AI-DI**, *right hand panels*), the SD of inter-breath intervals (**Fig 8AII-DII**), minute ventilation (**Fig 8E**), and apnoea frequency, duration and duration index (**Fig 8F I-III**). Clearly, ventilatory dysfunction worsened with progressive loss of Lkb1, but was most severe following *AMPKα1+α2* deletion.

**Figure 8.**
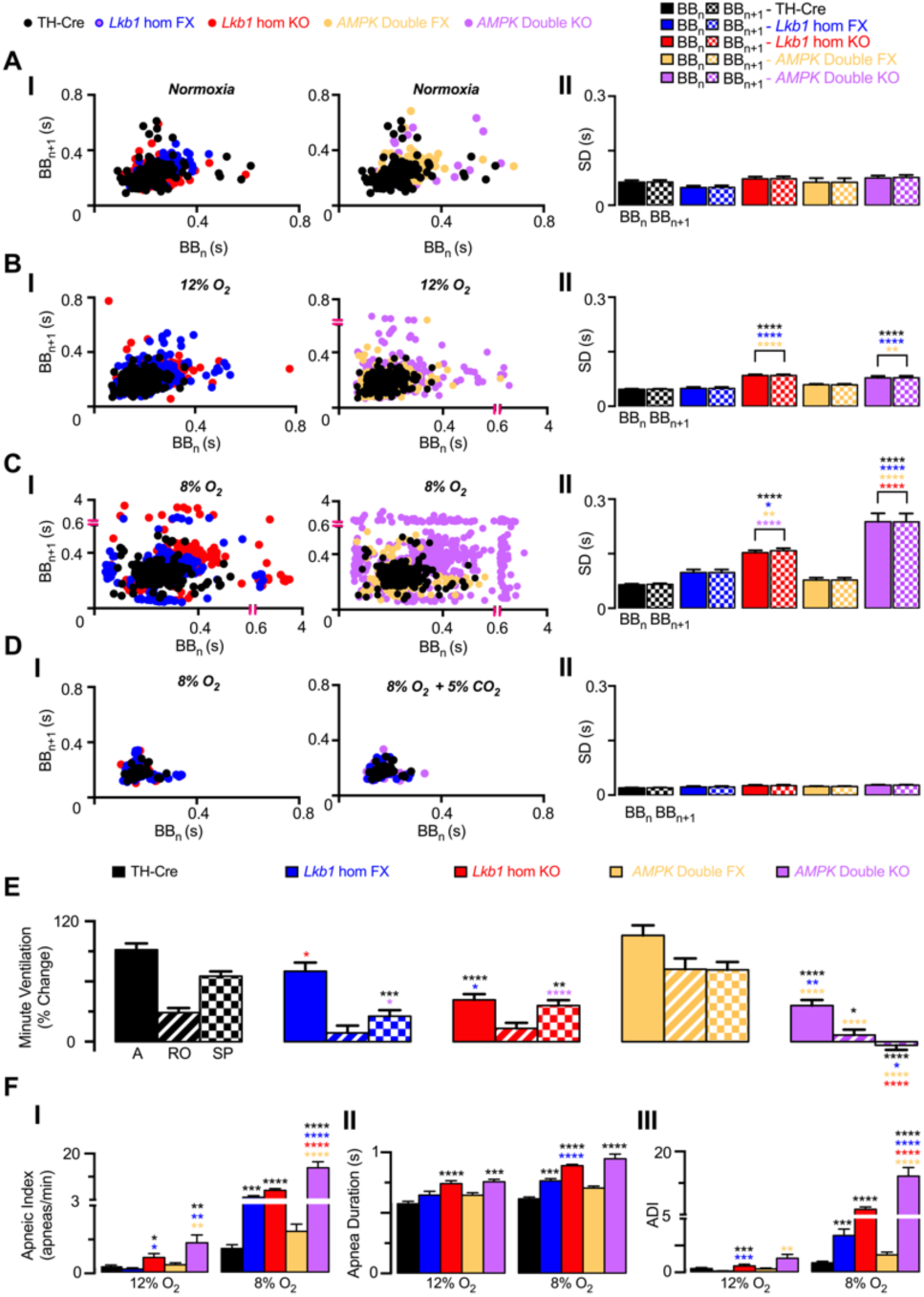
Unrestrained plethysmography reveals that respiratory dysfunction during hypoxia is less severe for conditional *Lkb1* deletion than for conditional *AMPK-*α*1+*α2 deletion in catecholaminergic cells. AI, *Lefthand panel* shows exemplar Poincaré plots of the inter-breath interval (BB_n_) versus subsequent interval (BB_n+1_) during normoxia for controls (TH-Cre, black, n=31), *Lkb1 homozygous* floxed (*Lkb1* hom FX, blue, n=14) and conditional *Lkb1* homozygous knockouts (*Lkb1* hom KO, red, n=22). *Right hand panel* shows comparable data for *AMPK-α1*+*α2* double floxed mice (*AMPK* hom FX, n=31) and conditional *AMPK-α1+α2* double knockouts (*AMPK* hom DKO, n=22). AII, Corresponding mean±SEM for the standard deviation (SD) of BB_n_ and BB_n+1_ for each genotype are shown for normoxia. BI, Poincaré plots of BB_n_ versus BB_n+1_ for mild hypoxia (12% O_2_). BII, Corresponding mean±SEM for the standard deviation (SD) of BB_n_ and BB_n+1_ for each genotype are shown for mild hypoxia (12% O_2_). CI, Poincaré plots of BB_n_ versus BB_n+1_ for severe hypoxia (8% O_2_). CII, Corresponding mean±SEM for the standard deviation (SD) of BB_n_ and BB_n+1_ for each genotype are shown for severe hypoxia (8% O_2_). DI, Poincaré plots of BB_n_ versus BB_n+1_ for hypercapnic hypoxia (8% O_2_ + 5% CO_2_). Corresponding mean±SEM for the standard deviation (SD) of BB_n_ and BB_n+1_ for each genotype are shown for hypercapnic hypoxia (8% O_2_ + 5% CO_2_). E, Comparison for all genotypes tested of mean±SEM for the % change of minute ventilation at the peak of the Augmenting Phase (∼30s), at ∼100s following Roll Off and during the plateau of the Sustained Phase (∼300s) during exposures to 8% O_2_. F, Comparison for all genotypes tested of the mean±SEM for (*I*) apnoeic index (per minute), (*II*) apnoea duration (s) and (*III*) apnoea-duration index (frequency × duration). *=p<0.05, **=p<0.01, *** =P<0.001, ****=p< 0.0001.

## 3. DISCUSSION

The present study demonstrates that deletion of *Lkb1* or *AMPKα1+α2*, but not *CaMKK2*, attenuates HVRs, precipitating hypoventilation and apnoea during hypoxia. The LKB1-AMPK signalling pathway is therefore critical to the delivery of ventilatory drive during hypoxia, and acts to oppose ventilatory depression, hypoventilation and apnoea. However, surprising differences between *Lkb1* and *AMPKα1+α2* knockouts were identified with respect to both HVRs and carotid body afferent fibre discharge, which provides further substantiation of our proposal [3] that AMPK facilitates HVRs downstream of peripheral chemoafferent input responses, most likely at the brainstem. Of particular note in this respect was the fact that LKB1 deletion attenuated HVRs and hypoxia-evoked increases in carotid body afferent discharge, while by contrast *AMPKα1+α2* deletion abolished HVRs but had little if any effect on carotid body afferent discharge during hypoxia [3]. Irrespective of the precise cellular mechanism, it is therefore clear that we have uncovered a split in the dependency on Lkb1 and AMPK of carotid body chemoafferent discharge during hypoxia on the one hand and HVRs on the other.

This view gained support from our studies on *Lkb1* floxed mice, which exhibit ∼90% global Lkb1 deficiency [9], but retain significantly greater capacity for basal and hypoxia-evoked carotid body afferent discharge than do *Lkb1* knockouts and yet still exhibit delayed hypoventilation and apnoea during severe hypoxia (8% O_2_). Intriguingly in this respect, deficits in minute ventilation were evident for *Lkb1* floxed mice during the sustained but not the augmenting phase. This is entirely consistent with an inhibitory effect downstream of chemoafferent input responses, if one accepts evidence supporting the view that increases in carotid body afferent discharge drive the augmenting phase of HVRs [15, 16] while direct modulation by hypoxia of brainstem respiratory networks maintains HVRs in the longer term [3-5, 16, 17]. Further support for this proposal may be derived from the fact that the order of severity for hypoxic ventilatory dysfunction increases progressively with the degree of Lkb1 deficiency (Fig. 5), but is most severe following *AMPK-α1+α2* deletion [3]. This is in spite of our finding that only *Lkb1* deletion attenuates carotid body afferent discharge, and suggests that the increased severity of ventilatory instability during sustained hypoxia is governed centrally by AMPK activity in catecholaminergic neurons.

Differences in outcome between genotypes provide yet further insights into the underlying circuit mechanisms governing HVRs. Hypomorphic *Lkb1* floxed mice exhibit marked attenuation of breathing frequency responses during severe hypoxia while tidal volume responses remained largely unaltered. AMPKα1/α2 deletion similarly attenuated only breathing frequency during mild hypoxia (12% O_2_), but blocked increases in both breathing frequency and tidal volume during severe hypoxia. Given that HVRs of mice hypomorphic for Lkb1 are less severely compromised than that of mice following *AMPKα1+α2* deletion, these outcomes suggest that the *P*O_2_-dependence for depression of breathing frequency is steeper than that for tidal volume in mice with deficiencies within the Lkb1-AMPK signalling pathway. AMPK may therefore contribute to each component of HVRs via divergent brainstem networks that lie downstream of carotid body afferent input responses, a point emphasised by the fact that *AMPKα1+α2* deletion is without effect on carotid body afferent discharge during hypoxia. That this may be the case gains further support from the generally held view that increases in breathing frequency during moderate to severe hypoxia are primarily driven by carotid body afferent input responses (see for example [21]).

Consistent with the aforementioned proposals, homozygous *Lkb1* deletion led to marked reductions in breathing frequency (excluding apnoeas) that were coupled with erratic “augmentation” of tidal volume responses during severe hypoxia. An explanation for this apparent increase in tidal volume was provided by the fact that *Lkb1* knockouts exhibited pronounced Cheyne-Stokes-like breathing (CSB) during hypoxia. That said, it is curious to note that CSB in *Lkb1* knockouts is accompanied by loss of carotid body afferent inputs given that hyperactivity of these peripheral chemoreceptors has been identified as the cause of CSB when associated with heart failure [25]. One possible explanation for this contrary outcome could be that *Lkb1* deletion alone attenuated basal afferent discharge from the carotid body and virtually abolished increases in chemoafferent discharge during hypoxia and hypercapnia, actions that could serve to attenuate any inhibitory interactions between peripheral chemoreceptors and brainstem respiratory networks [15, 24]. That this might be the case is indicated by the fact that the peak of the sustained phase of the hypercapnic ventilatory response of *Lkb1* knockouts remained unaltered despite the reduction of afferent input responses. In other words, increases in controller gain within the central respiratory network could trigger CSB by enhancing the sensitivity to, and thus the degree of activation of central CO_2_-sensing neurons during hypercapnia [15] consequent to hypoventilation during hypoxia, leading to periodic augmentation of tidal volume in *Lkb1* knockouts alone. Consistent with this view, others have proposed that CSB may be caused by enhanced hypercapnic ventilatory responses driven by instability within respiratory networks consequent to either augmented chemoreflex gain, prolonged feedback delay [26] and/or enhanced central controller gain [27]. In short, apnoeic intervals may well be countered earlier in *Lkb1* knockouts through augmented central hypercapnic ventilatory responses consequent to reductions in basal and evoked carotid body afferent discharge frequencies [15, 24], resulting in CSB during periods of hypoxia-evoked reductions in breathing frequency. The more extreme patterns of non-rhythmic (ataxic) ventilation observed for *AMPKα1/α2* knockouts [3] may thus be avoided. While less likely it is also conceivable that retention by *Lkb1* knockouts of greater capacity for rhythmic ventilation during hypoxia could be conferred by residual allosteric AMPK activation by AMP, because this occurs independent of Lkb1 [1] and any fall in cellular ATP supply during hypoxia would be associated with not only ADP accumulation but consequent increases in the AMP:ATP ratio via the adenylate kinase reaction. This may in its own right be sufficient to maintain oscillating central respiratory drive in a manner periodically triggered once a given severity of central hypoxia is breached. That said, if sighs are triggered by hypoxia at a given threshold [22, 28], central hypoxia is likely no more severe for *Lkb1* when compared to *AMPKα1+α2* knockouts because: (1) A similar frequency of sighs is observed during hypoxia for each of these genotypes; (2) Apnoeas are shorter and less frequent for *Lkb1* when compared to *AMPKα1+α2* knockouts; (3) Only *Lkb1* knockouts exhibit CSB between apnoeas, which would periodically raise O_2_ supply. Therefore, unless loss of carotid body afferent input responses in *Lkb1* knockouts alters functional hyperaemia at the level of the brainstem and via neuronal pathways independent of those mediating HVRs, it seems most likely that differences in outcome between genotypes arise from means other than variations in severity of brainstem hypoxia. However, for this question to be answered conclusively we would need to identify the precise hypoxia-responsive brainstem region(s) affected by our gene deletion strategies and obtain direct, region-specific measurements of local *P*O_2_.

While the aforementioned findings run counter to the view that increased afferent discharge from carotid body to brainstem alone determines the ventilatory response to a fall in arterial *P*O_2_ they do provide substantial support for an alternative yet inclusive perspective, namely that HVRs are determined by the coordinated action of the carotid body and an hypoxia-responsive circuit within the brainstem [3, 5, 16, 17]. To date little emphasis has been placed on the role of hypoxia-sensing downstream, at the level of the brainstem perhaps because HVRs are so effectively abolished by resection of the carotid sinus nerve in humans [29]. And yet brainstem hypoxia induces an HVR when in receipt of normoxic carotid body afferent inputs [17], and directly activates subsets of catecholaminergic neurons within the brainstem nucleus of the solitary tract [30] and rostral ventrolateral medulla [5, 31, 32] in a manner that may be supported by direct activation by hypoxia of ATP/lactate release from brainstem astrocytes [33, 34]. Moreover and consistent with the fact that our gene deletion strategy targeted catecholaminergic neurons, ectopic expression aside, extensive investigations have demonstrated that following carotid body resection, hypoxia-responsive catecholaminergic neurons of the caudal brainstem may underpin partial recovery of the HVR in a variety of animal models [5]. Accordingly, dysfunction of these neurons has been shown to underpin hypoventilation and apnoea associated with Rett syndrome, which is exacerbated during hypoxia [35].

Insights into the mechanisms that determine afferent discharge from the carotid body may also be garnered from our observation that *Lkb1* but not *AMPKα1+α2* deletion markedly attenuated basal afferent fibre discharge and blocked increases in afferent discharge during hypercapnia, given that increases in afferent discharge during hypercapnia are triggered by membrane depolarisation consequent to hypercapnic acidosis and in a manner not directly influenced by reductions in mitochondrial oxidative phosphorylation or deficits in ATP supply [36]. This suggests that while *Lkb1* contributes to the maintenance of carotid body afferent discharge, it does not necessarily support type I cell oxygen-sensing *per se*. We must, however, add a note of caution here, because this assay does not directly distinguish between actions on the type I cells and tyrosine hydroxylase expressing glossopharyngeal nerves. Nevertheless, we can conclude that Lkb1 expression is somehow necessary for chemoafferent outflow from the carotid body. That said, the reduced hypercapnic chemoafferent responses could alternatively be due to the removal of CO_2_-O_2_ stimulus interaction [37, 38]. Either way, our findings suggest that Lkb1 determines, independent of AMPK, carotid body chemoafferent discharge [39]. Lkb1 could conceivable contribute to developmental expansion of carotid body type I cells, or to the regulation of glucose homeostasis [40, 41] and mitochondrial function [42, 43], either directly or in a manner supported by constitutive phosphorylation of one or more of the 12 AMPK-related kinases [44]. In short, a role for Lkb1 in the carotid body that is independent of the modification in AMPK activity is intriguing, but further, extensive investigations will be required to determine the precise mechanism(s) involved.

## 4. CONCLUSION

Lkb1 and AMPK provide hierarchical control of the chemo-sensory respiratory network. Firstly, Lkb1 appears to determine, independent of AMPK, a set-point about which carotid body afferent input responses are evoked during hypoxia and hypercapnia, rather than contributing to oxygen-sensing *per se*. Thereafter the Lkb1-AMPK signalling pathway likely governs, through the capacity for AMPK activation by increases in AM(D)P/ATP ratio and Lkb1-dependent phosphorylation, coincidence detection and thus signal integration within a hypoxia-responsive circuit downstream of the carotid body, that encompasses, at the very least, the nucleus of the solitary tract and ventrolateral medulla [3]. Afferent input responses and brainstem hypoxia could thereby determine, each in part, the set-point about which AMPK and thus the brainstem respiratory networks are activated during hypoxia. Subsequently, AMPK-dependent modulation of cellular metabolism[1], ion channels [23, 45] and thus neuronal activities [46, 47] may facilitate efferent output and increases in ventilatory drive during hypoxia. It is therefore conceivable that Lkb1 and/or AMPK deficiency may contribute to Cheyne-Stokes breathing [26] and/or sleep disordered breathing associated with, for example, heart failure [25], metabolic syndrome-related disorders[48] and ascent to altitude [49].

## 5. METHODS

Experiments were carried out as described previously[3], were approved by local ethical review committees, the University Director of Veterinary Services and the Home Office (Science, UK), and complied with the regulations of the United Kingdom Animals (Scientific Procedures) Act of 1986.

### 5.1 Breeding of mice, genotyping and single cell PCR

Standard approaches were used. All mice studied were between 3-12 months of age. Both males and females were studied.

For *Lkb1* deletion we used mice with exons 5-7 of the *Lkb1* gene (STK11) replaced by a cDNA cassette encoding equivalent exon sequences, and exon 4 and the cDNA cassette flanked by loxP sequences (*Lkb1*^*fl/fl*^). These mice were crossed with transgenic mice expressing Cre-recombinase under the tyrosine hydroxylase promoter (Th-IRES-Cre; EM:00254)[12]. Wild type or floxed *Lkb1* alleles were detected using two primers, p200, 5’-CCAGCCTTCTGACTCTCAGG-3’ and p201, 5’-GTAGGTATTCCAGGCCGTCA-3’. For the detection of CRE recombinase we employed: TH3, 5’-CTTTCCTTCCTTTATTGAGAT-3’, TH5, 5’-CACCCTGACCCAAGCACT-3’ and Cre-UD, 5’-GATACCTGGCCTGGTCTCG-3’. As *Lkb1*^*fl/fl*^ mice are hypomorphic, exhibiting 5-10 fold lower LKB1 expression than *Lkb1*^+/+^ littermates [9], we used as controls mice that express Cre via the tyrosine hydroxylase promoter (TH-Cre).

For deletion of the gene that encodes CaMKK2 (*CaMKK2*) wild type alleles were detected using two primers, KKBeta1, 5’CAGCACTCAGCTCCAATCAA3’, and KKBeta2, 5’GCCACCTATTGCC TTGTTTG3’. The PCR protocol used for all genotype primers was: 92°C for 5min, 92°C for 45s, 56°C for 45s, 72°C for 60s, and 72°C for 7min for 35 cycles and then 4°C as the holding temperature. 15 µl samples were run on 2% agarose gels with 10 µl SYBR®Safe DNA Gel Stain (Invitrogen) in TBE buffer against a 100 bp DNA ladder (GeneRuler™, Fermentas) using a Model 200/2.0 Power Supply (Bio-Rad). Gels were imaged using a Genius Bio Imaging System and GeneSnap software (Syngene).

We also used conditional deletion of the genes for the *AMPKα1* and *α2* subunits, utilising mice in which the sequence encoding the catalytic site of both of the α subunits was flanked by loxP sequences (*α1*^*flx*^ *and α2*^*flx*^ [10]). We used two primers for each AMPK catalytic subunit: α1-forward: 5’ TATTGCTGCCATTAGGCTAC 3’, α1-reverse: 5’ GACCTGACAGAATAGGATATGCCCAACCTC 3’; α2-forward 5’ GCTTAGCACGTTACCCTGGATGG 3’, α2-reverse: 5’ GTTATCAGCCCAACTAATTACAC 3’. To direct *AMPK* deletion to identified oxygen-sensing cells of the carotid body and brainstem, these were crossed with TH-Cre mice as above. We detected the presence of wild-type or floxed *AMPK* alleles *and* Cre-recombinase by PCR. 15 µl samples were run on 2% agarose gels and imaged as described above.

### 5.2 Single-cell end-point PCR

Carotid bodies were incubated at 37°C for 25-30 min in isolation medium consisting of: 0.125mg/ml Trypsin (Sigma), 2.5mg/ml collagenase Type 1 (Worthington) made up in low Ca^2+^/low Mg^2+^ HBSS. During this incubation the carotid bodies were separated from the associated patch of artery. The carotid bodies were then transferred to low Ca^2+^/low Mg^2+^ HBSS containing trypsin inhibitor (0.5mg/ml) for 5min at room temperature, and then to 2ml of pre-equilibrated (95% air, 5% CO_2_, 37°C) growth medium (F-12 Ham nutrient mix, 10% fetal bovine serum, 1% penicillin/streptomycin). The medium containing the carotid bodies was centrifuged and the pellet re-suspended in 100µl of growth medium. Carotid bodies were then disrupted by triturating using fire polished Pasteur pipettes.

RNA was extracted using the High Pure RNA Tissue Kit (Roche) following the manufacturer’s guidelines and the concentration determined using the Nanodrop 1000 spectrophotometer (ThermoScientific). cDNA synthesis was carried out using the Transcriptor High Fidelity cDNA synthesis Kit (Roche) following the manufacturers’ instructions. Amplification of cDNA isolated from different individual cells was run in parallel with negative and positive controls using an initial denaturing step at 94°C for 5min and then denaturing at 94°C for 30s, annealing at 60°C for 45s, and extension for 60s at 72°C with a final 7min extension at 72°C. Initially 15 cycles were performed, followed by reaction and dilution for a further 38 cycles. To detect tyrosine hydroxylase, primers obtained from Qiagen (Quantitect Primer Assay, QT00101962) were used with an expected band length of 96bp. For the detection of Lkb1 two primers were used, *forward* and *reverse*, to generate an expected band length of 92bp.

Negative controls included control cell aspirants, lacking reverse transcriptase, aspiration of extracellular medium and PCR controls; these produced no detectable amplicons, ruling out genomic or other contamination. 15µl samples and a 100bp DNA ladder (GeneRuler™, Fermentas) were run on 2% agarose gels with SYBR^®^Safe DNA Gel Stain (Invitrogen). Gels were imaged using a Genius Bio Imaging System and GeneSnap software (Syngene). Positive controls were from samples rich in adrenomedullary chromaffin cells, dissected from adrenal glands of C57/Bl6 mice. RNA was extracted using the High Pure RNA Tissue Kit (Roche) following the manufacturer’s guidelines and the concentration determined using the Nanodrop 1000 spectrophotometer (ThermoScientific). cDNA synthesis was carried out using the Transcriptor High Fidelity cDNA kit (Roche) following manufacturers instructions.

### 5.3 Quantitative RT-PCR

RNA from adrenal glands was extracted, quantified and reverse transcribed as described above. For qPCR analysis, 2.5 µl of cDNA in RNase free water was made up to 25 µl with FastStart Universal SYBR Green Master (ROX, 12.5 µl, Roche), Ultra Pure Water (8 µl, SIGMA) and forward and reverse primers for Lkb1. The sample was then centrifuged and 25 µl added to a MicroAmp™ Fast Optical 96-Well Reaction Plate (Greiner bio-one), the reaction plate sealed with an optical adhesive cover (Applied Biosystems) and the plate centrifuged. The reaction was then run on a sequence detection system (Applied Biosystems) using AmpliTaq Fast DNA Polymerase, with a 2min initial step at 50°C, followed by a 10min step at 95°C, then a 15s step at 95°C which was repeated 40 times. Then a dissociation stage with a 15s step at 95°C followed by a 20s at 60°C and a 15s step at 95°C. Negative controls included control cell aspirants for which no reverse transcriptase was added, and aspiration of extracellular medium and PCR controls. None of the controls produced any detectable amplicon, ruling out genomic or other contamination.

### 5.4 Plethysmography

We used unrestrained whole-body plethysmography, incorporating a Halcyon™ low noise pneumatochograph coupled to FinePointe acquisition and analysis software with a sampling frequency of 1kHz (Buxco Research Systems, UK). All quoted values for HVR were derived from apnoea-free periods of ventilation. Any unreliable and erratic respiratory waveforms recorded during gross un-ventilatory related body movements, i.e. sniffing and grooming, were avoided for measurements. Additionally, a rejection algorithm that was built into the plethysmography system (Buxco Electronics Inc.) identified periods of motion-induced-artefacts for omission. The patented Halycon™ low noise pneumotachograph (Buxco Electronics Inc.) reduces disturbances caused by air currents from outside the chambers (i.e. fans, closing doors, air conditioners, etc.), which can disrupt or overwhelm the ventilatory airflows within the chamber.

Mice were placed in the plethysmography chamber for a 10-20 min acclimation period under normoxia (room air) to establish a period of quiet and reliable breathing for baseline-ventilation levels (this is also indicated by a measured rejection index of 0 by the FinePointe Acquisition and Analysis Software). Mice were then exposed to hypoxia (12% or 8% O_2_, with 0.05% CO_2_, balanced with N_2_), hypoxia+hypercapnia (8% O_2_, 5% CO_2_, balanced with N_2_) or hypercapnia (21% O_2_, 5% CO_2_, balanced with N_2_) for 5min. Medical grade gas mixtures were chosen by switching a gas tap. The time for evacuation of the dead space and complete exchange of gas within the plethysmography chamber was 30s. The duration of exposure to hypoxia quoted was the actual duration of hypoxia. Apnoea was defined as cessations of breathing greater than the average duration, including interval, of 2 successive breaths (600ms) during normoxia, with a detection threshold of 0.25 mmHg (SD of noise). Breathing variability was assessed by Poincaré plots and by calculating the SD of inter-breath (BB) intervals. The breathing frequency, tidal volume, and minute ventilation as derived by the FinePointe Software were also analysed for control and knockout mice. These parameters were measured as mean values taken over a 2s breathing period and not on a breath-to-breath basis. The changes in breathing frequency, tidal volume, and minute ventilation during hypoxia and/or hypercapnia were analysed as the percentage change from normoxia respective to each individual mouse. The peak of the augmenting phase (A) was calculated from the peak value between 20-40s of the hypoxic and/or hypercapnic exposure that coincides with the peak of the rising phase. The roll off period was calculated as the lowest value between 60-140s of exposure and the sustained phase was calculated from the last 20s in the plateaued phase. A large time range was required for selection of these points as experiments were performed on unrestrained and awake animals and periods of no movement, sniffing, or grooming, were only considered.

Apnoeas were excluded from all stated measures (mean±SEM) of breathing frequency, tidal volume and minute ventilation, i.e., all quoted values were derived from apnoea-free periods of ventilation.

### 5.5 Isolated carotid body

Methods for single fibre chemoafferent activity were adapted from those described previously[14, 50]. Plots of firing frequency versus superfusate *p*O_2_ were fitted by non-linear regression (GraphPad Prism 6).

Single fibre chemoafferent activity was amplified and filtered and recorded using a 1401 interface running Spike 2 software (Cambridge Electronic Design). Single- or few-fibre chemoafferent recordings were made from carotid bifurcations held in a small volume tissue bath, and superfused (36-37°C) with gassed (95% O_2_ and 5% CO_2_), bicarbonate-buffered saline solution (composition (mM): 125 NaCl, 3 KCl, 1.25 NaH_2_PO_4_, 5 Na_2_SO_4_, 1.3 MgSO_4_, 24 NaHCO_3_, 2.4 CaCl_2_). A standard O_2_ electrode (ISO2; World Precision Instruments) was placed in the superfusate system at the point of entry to the recording chamber in order to continuously record the superfusate *P*O_2_. Flow meters with high precision valves (Cole Palmer Instruments) were used to equilibrate the superfusate with a desired gas mixture. Basal single fibre activity was monitored at a superfusate PO_2_ of 200mmHg and a PCO_2_ of 40mmHg. This PO_2_ is slightly lower than that previously used for the rat carotid body[51] to take in account the smaller size of this organ in the mouse (and thus a smaller diffusion distance). At this superfusate PO_2_, the basal frequency in TH-Cre single fibres (Figure 1) was consistent with that reported *in vivo* in other rodents[52] and so we interpret this PO_2_ to have not been excessively hyperoxic.

To induce responses to hypoxia, the superfusate *P*O_2_ was slowly reduced to a minimum of 40 mmHg or was reversed prior to this when the chemoafferent response had stabilised or had begun to diminish. The single fibre chemoafferent discharge frequency was plotted against the superfusate *P*O_2_ over a desired range of superfusate *P*O_2_ values. To produce the hypoxic response curves, the data points were fitted to an exponential decay curve with offset, as shown below:

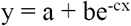

For the above equation, y is the single fibre discharge frequency in Hz, x is the superfusate *P*O_2_ in mmHg, a is the discharge frequency as the *P*O_2_ tends to infinity (offset), b is the discharge frequency when the *P*O_2_ is 0 mmHg (minus the offset) and c is the exponential rate constant.

### 5.6 Statistical analysis

Statistical comparison was completed using GraphPad Prism 6 for the following: *Afferent discharge*, single or 2 factor ANOVA with Bonferroni Dunn post hoc analysis; *Plethysmography*, one-way ANOVA with Bonferroni multiple comparison’s test; P<0.05 was considered significant.

## AUTHORS’ CONTRIBUTIONS

A.M.E. wrote the manuscript and made Figures 3, 6 and 7. A.D.M made Figures 1, 2, 4 and 8 and supplementary Figures 1-4. S.M. made Figure 5. A.M.E. and O.A.O developed the conditional Lkb1 knockout mice. A.M.E., A.D.M. and S.M. developed the conditional AMPK knockout mice, and performed genotyping. M.F. and B.V. developed the AMPK floxed mice. A.D.M. designed and validated primers. A.D.M. performed single cell PCR. A.M.D, S.M., and A.M.E. performed plethysmography on LKB1 and/or AMPK knockouts. A.D.M., S.M., and A.M.E. analysed respiratory data. A.P.H. and P.K. performed afferent discharge blind and under subcontract at the University of Birmingham. C.N.W. developed the murine carotid body type I cell isolation, and trained M.L.D., who carried out the type I cell isolation in support of this study.

## ACKNOWLEDGEMENTS

This work was primarily funded by the Wellcome Trust (WT081195MA). A.M.E. thanks Professor D. Grahame Hardie for his continued guidance, support and for providing the Lkb1 floxed mice. We would also like to thank Professor Michael J. Shipston and Helene Widmer for their kind help and assistance with the single cell PCR shown in the supplementary information.

## REFERENCES

1. Ross F.A., MacKintosh C. & Hardie D.G. AMP-activated protein kinase: a cellular energy sensor that comes in 12 flavours. FEBS J. 83, 2987–3001 (2016).

2. Gowans G.J., Hawley S.A., Ross F.A. & Hardie D.G. AMP is a true physiological regulator of AMP-activated protein kinase by both allosteric activation and enhancing net phosphorylation. Cell Metab. 18, 556–566 (2013).

3. Mahmoud A.D., Lewis S., Juricic L., Udoh U.A., Hartmann S., Jansen M.A., Ogunbayo O.A., Puggioni P., Holmes A.P., Kumar P., Navarro-Dorado J., Foretz M., Viollet B., Dutia M.B., Marshall I. & Evans A.M. AMPK Deficiency Blocks the Hypoxic Ventilatory Response and Thus Precipitates Hypoventilation and Apnea. Am. J. Respir. Crit. Care Med. 193, 1032–1043 (2016).

4. Wilson R.J. & Teppema L.J. Integration of Central and Peripheral Respiratory Chemoreflexes. Compr. Physiol. 6, 1005–1041 (2016).

5. Teppema L.J. & Dahan A. The ventilatory response to hypoxia in mammals: mechanisms, measurement, and analysis. Physiol. Rev. 90, 675–754 (2010).

6. Guyenet P.G. Neural structures that mediate sympathoexcitation during hypoxia. Resp. Physiol. 121, 147–162 (2000).

7. Smith J.C., Abdala A.P., Borgmann A., Rybak I.A. & Paton J.F. Brainstem respiratory networks: building blocks and microcircuits. Trends. Neurosci. 36, 152–162 (2013).

8. Nurse C.A. Synaptic and paracrine mechanisms at carotid body arterial chemoreceptors. J. Physiol. 592, 3419–3426 (2014).

9. Sakamoto K., McCarthy A., Smith D., Green K.A., Hardie D.G., Ashworth A. & Alessi D.R. Deficiency of LKB1 in skeletal muscle prevents AMPK activation and glucose uptake during contraction. EMBO J. 24, 1810–1820 (2005).

10. Lantier L., Fentz J., Mounier R., Leclerc J., Treebak J.T., Pehmoller C., Sanz N., Sakakibara I., Saint-Amand E., Rimbaud S., Maire P., Marette A., Ventura-Clapier R., Ferry A., Wojtaszewski J.F., Foretz M. & Viollet B. AMPK controls exercise endurance, mitochondrial oxidative capacity, and skeletal muscle integrity. FASEB J, 28, 3211–3224 (2014).

11. Hirooka Y., Polson J.W., Potts P.D. & Dampney R.A. Hypoxia-induced Fos expression in neurons projecting to the pressor region in the rostral ventrolateral medulla. Neuroscience 80, 1209–1224 (1997).

12. Lindeberg J., Usoskin D., Bengtsson H., Gustafsson A., Kylberg A., Soderstrom S. & Ebendal T. Transgenic expression of Cre recombinase from the tyrosine hydroxylase locus. Genesis 40, 67–73 (2004).

13. Anderson K.A., Ribar T.J., Lin F., Noeldner P.K., Green M.F., Muehlbauer M.J., Witters L.A., Kemp B.E. & Means A.R. Hypothalamic CaMKK2 contributes to the regulation of energy balance. Cell Metab. 7, 377–388 (2008).

14. Mahmoud A.D., Lewis S., Juricic L., Udoh U.A., Hartmann S., Jansen M.A., Ogunbayo O.A., Puggioni P., Holmes A.P., Kumar P., Navarro-Dorado J., Foretz M., Viollet B., Dutia M.B., Marshall I. & Evans A.M. AMP-activated Protein Kinase Deficiency Blocks the Hypoxic Ventilatory Response and Thus Precipitates Hypoventilation and Apnea. Am. J. Respir. Crit. Care Med. 193, 1032–1043 (2016).

15. Day T.A. & Wilson R.J. Brainstem PCO2 modulates phrenic responses to specific carotid body hypoxia in an in situ dual perfused rat preparation. J. Physiol. 578, 843–857 (2007).

16. Smith C.A., Engwall M.J., Dempsey J.A. & Bisgard G.E. Effects of specific carotid body and brain hypoxia on respiratory muscle control in the awake goat. J. Physiol. 460, 623–640 (1993).

17. Curran A.K., Rodman J.R., Eastwood P.R., Henderson K.S., Dempsey J.A. & Smith C.A. Ventilatory responses to specific CNS hypoxia in sleeping dogs. J. Appl. Physiol. 88, 1840–1852 (2000).

18. Chapman R.W., Santiago T.V. & Edelman N.H. Effects of graded reduction of brain blood flow on ventilation in unanesthetized goats. J. Appl. Physiol. Respir. Environ. Exerc. Physiol. 47, 104–111 (1979:).

19. Martin-Body R.L., Robson G.J. & Sinclair J.D. Restoration of hypoxic respiratory responses in the awake rat after carotid body denervation by sinus nerve section. J. Physiol. 380, 61–73 (1986).

20. Hodson E.J., Nicholls L.G., Turner P.J., Llyr R., Fielding J.W., Douglas G., Ratnayaka I., Robbins P.A., Pugh C.W., Buckler K.J., Ratcliffe P.J. & Bishop T. Regulation of ventilatory sensitivity and carotid body proliferation in hypoxia by the PHD2/HIF-2 pathway. J. Physiol. 594, 1179–1195 (2016).

21. Martin-Body R.L., Robson G.J. & Sinclair J.D. Respiratory effects of sectioning the carotid sinus glossopharyngeal and abdominal vagal nerves in the awake rat. J Physiol 1361, 35–45 (1985).

22. Li P., Janczewski W.A., Yackle K., Kam K., Pagliardini S., Krasnow M.A. & Feldman J.L. The peptidergic control circuit for sighing. Nature 530, 293–297 (2016).

23. Ross F.A., Rafferty J.N., Dallas M.L., Ogunbayo O., Ikematsu N., McClafferty H., Tian L., Widmer H., Rowe I.C., Wyatt C.N., Shipston M.J., Peers C., Hardie D.G. & Evans A.M. Selective Expression in Carotid Body Type I Cells of a Single Splice Variant of the Large Conductance Calcium- and Voltage-activated Potassium Channel Confers Regulation by AMP-activated Protein Kinase. J. Biol. Chem. 286, 11929–11936 (2011).

24. Blain G.M., Smith C.A., Henderson K.S. & Dempsey J.A. Peripheral chemoreceptors determine the respiratory sensitivity of central chemoreceptors to CO(2). J Physiol 588, 2455–2471 (2010).

25. Ponikowski P., Chua T.P., Anker S.D., Francis D.P., Doehner W., Banasiak W., Poole-Wilson P.A., Piepoli M.F. & Coats A.J. Peripheral chemoreceptor hypersensitivity: an ominous sign in patients with chronic heart failure. Circulation 104, 544–549 (2001).

26. Hall M.J., Xie A., Rutherford R., Ando S., Floras J.S. & Bradley T.D. Cycle length of periodic breathing in patients with and without heart failure. Am. J. Respir. Crit. Care Med. 154, 376–381 (1996).

27. Topor Z.L., Vasilakos K., Younes M. & Remmers J.E. Model based analysis of sleep disordered breathing in congestive heart failure. Resp. Physiol. Neurobiol. 155, 82–92 (2007).

28. Bell H.J. & Azubike E, Haouzi P. The “other” respiratory effect of opioids: suppression of spontaneous augmented (“sigh”) breaths. J. Appl. Physiol. 111, 1296–1303 (2011).

29. Wade J.G., Larson C.P. Jr., Hickey R.F., Ehrenfeld W.K. & Severinghaus J.W. Effect of carotid endarterectomy on carotid chemoreceptor and baroreceptor function in man. New England J. Med. 282, 823–829 (1970).

30. Pascual O., Morin-Surun M.P., Barna B., Denavit-Saubie M., Pequignot J.M. & Champagnat J. Progesterone reverses the neuronal responses to hypoxia in rat nucleus tractus solitarius in vitro. J. Physiol. 544, 511–520 (2002).

31. Nolan P.C. & Waldrop T.G. In vivo and in vitro responses of neurons in the ventrolateral medulla to hypoxia. Brain Res. 630, 101–114 (1993).

32. Sun M.K. & Reis D.J. Differential responses of barosensitive neurons of rostral ventrolateral medulla to hypoxia in rats. Brain Res. 609, 333–337 (1993).

33. Angelova P.R., Kasymov V., Christie I., Sheikhbahaei S., Turovsky E., Marina N., Korsak A., Zwicker J., Teschemacher A.G., Ackland G.L., Funk G.D., Kasparov S., Abramov A.Y. & Gourine A.V. Functional Oxygen Sensitivity of Astrocytes. J. Neurosci. 35, 10460–10473 (2015).

34. Magistretti P.J. & Allaman I. Lactate in the brain: from metabolic end-product to signalling molecule. Nat. Rev. Neurosci. 19, 235–249 (2018).

35. Roux J.C. & Villard L. Biogenic amines in Rett syndrome: the usual suspects. Behavior Genetics 40, 59–75 (2010).

36. Mulligan E. & Lahiri S. Separation of carotid body chemoreceptor responses to O2 and CO2 by oligomycin and by antimycin A. Am. J. Physiol. 242, C200–206 (1982).

37. Dasso L.L., Buckler K.J., Vaughan-Jones R.D. Interactions between hypoxia and hypercapnic acidosis on calcium signaling in carotid body type I cells. Am. J. Physiol. 279, L36–42 (2000).

38. Pepper D.R., Landauer R.C. & Kumar P. Postnatal development of CO2-O2 interaction in the rat carotid body in vitro. J. Physiol. 485, 531–541 (1995).

39. Murali S. & Nurse C.A. Purinergic signalling mediates bidirectional crosstalk between chemoreceptor type I and glial-like type II cells of the rat carotid body. J. Physiol. 594, 391–406 (2016).

40. Shaw R.J., Lamia K.A., Vasquez D., Koo S.H., Bardeesy N., Depinho R.A., Montminy M. & Cantley L.C. The kinase LKB1 mediates glucose homeostasis in liver and therapeutic effects of metformin. Science 310, 1642–1646 (2005).

41. Patel K., Foretz M., Marion A., Campbell D.G., Gourlay R., Boudaba N., Tournier E., Titchenell P., Peggie M., Deak M., Wan M., Kaestner K.H., Goransson O., Viollet B., Gray N.S., Birnbaum M.J., Sutherland C. & Sakamoto K. The LKB1-salt-inducible kinase pathway functions as a key gluconeogenic suppressor in the liver. Nature Comm. 5, 4535 (2014).

42. Gan B., Hu J., Jiang S., Liu Y., Sahin E., Zhuang L., Fletcher-Sananikone E., Colla S., Wang Y.A., Chin L., Depinho R.A. Lkb1 regulates quiescence and metabolic homeostasis of haematopoietic stem cells. Nature 468, 701–704 (2010).

43. Swisa A., Granot Z., Tamarina N., Sayers S., Bardeesy N., Philipson L., Hodson D.J., Wikstrom J.D., Rutter G.A., Leibowitz G., Glaser B. & Dor Y. Loss of Liver Kinase B1 (LKB1) in Beta Cells Enhances Glucose-stimulated Insulin Secretion Despite Profound Mitochondrial Defects. J. Biol. Chem. 290, 20934–20946 (2015).

44. Lizcano J.M., Goransson O., Toth R., Deak M., Morrice N.A., Boudeau J., Hawley S.A., Udd L., Makela T.P., Hardie D.G., Alessi D.R. LKB1 is a master kinase that activates 13 kinases of the AMPK subfamily, including MARK/PAR-1. EMBO J. 23, 833–843 (2004).

45. Ikematsu N., Dallas M.L., Ross F.A., Lewis R.W., Rafferty J.N., David J.A., Suman R., Peers C., Hardie D.G. & Evans A.M. Phosphorylation of the voltage-gated potassium channel Kv2.1 by AMP-activated protein kinase regulates membrane excitability. PNAS 108, 18132–18137 (2011).

46. Lipton A.J., Johnson M.A., Macdonald T., Lieberman M.W., Gozal D. & Gaston B. Snitrosothiols signal the ventilatory response to hypoxia. Nature 413, 171–174 (2001).

47. Murphy B.A., Fakira K.A., Song Z., Beuve A., Routh V.H. AMP-activated protein kinase and nitric oxide regulate the glucose sensitivity of ventromedial hypothalamic glucose-inhibited neurons. Am. J. Physiol. 297, C750–758 (2009).

48. Chau E.H., Lam D., Wong J., Mokhlesi B. & Chung F. Obesity hypoventilation syndrome: a review of epidemiology, pathophysiology, and perioperative considerations. Anesthesiology 117, 188–205 (2012).

49. Ainslie P.N., Lucas S.J. & Burgess K.R. Breathing and sleep at high altitude. Respir. Physiol. Neurobiol. 188, 233–256 (2013).

50. Wyatt C.N., Mustard K.J., Pearson S.A., Dallas M.L., Atkinson L., Kumar P., Peers C., Hardie D.G. & Evans A.M. AMP-activated protein kinase mediates carotid body excitation by hypoxia. J. Biol. Chem. 282, 8092–8098 (2007).

51. Holmes A.P., Turner P.J., Buckler K.J. & Kumar P. Moderate inhibition of mitochondrial function augments carotid body hypoxic sensitivity. Pflugers Arch. 468, 143–155 (2016).

52. Vidruk E.H., Olson E.B. Jr., Ling L. & Mitchell G.S. Responses of single-unit carotid body chemoreceptors in adult rats. J. Physiol. 531, 165–170 (2001).

